# *Vibrio* pore-forming leukocidin activates pyroptotic cell death via the NLRP3 inflammasome

**DOI:** 10.1101/816553

**Authors:** Noam Baram, Hadar Cohen, Liat Edry-Botzer, Dor Salomon, Motti Gerlic

## Abstract

Cell death mechanisms are central to combat infectious microbes and to drive pathological inflammation. One such mechanism, the inflammasome, controls infection through either activation of caspase-1 and the subsequent secretion of the mature pro-inflammatory cytokine, interleukin 1β (IL-1β), or by stopping the dissemination of intracellular pathogens by inducing pyroptotic cell death in infected cells. Hemolysins, which are pore-forming toxins (PFTs), target the host cell plasma membrane by producing pores with different diameters. These pores alter the permeability of the target membrane, often leading to cell death. We previously discovered a functional and potent pore-forming, leukocidin domain-containing hemolysin produced by the Gram-negative marine bacterium *Vibrio proteolyticus* (*V. proteolyticus*), termed VPRH. Although leukocidin domains are found in other known PFTs, VPRH constitutes a distinct, understudied class within the leukocidin superfamily. Since PTFs of other pathogens were shown to induce cell death by activating the inflammasome pathway, we hypothesized that VPRH-induced cell death is mediated by direct activation of the inflammasome in mammalian immune host cells. Indeed, we found that VPRH induced a two-step cell death in primary macrophages. The first, a rapid step, was mediated by activating the NLRP3 inflammasome, leading to caspase-1 activation and GSDMD cleavage that resulted in IL-1β secretion and pyroptotic cell death. The second step was independent of the inflammasome; however, its mechanism remains unknown. This study sets the foundation for better understanding the immunological consequences of inflammasome activation by a new leukocidin class of toxins.

## INTRODUCTION

Pathogenic bacteria often produce and secrete toxins to manipulate their hosts or to defy predators. A widespread and abundant class of virulence factors are pore-forming toxins (PFTs) [1]. Pore-forming hemolysins oligomerize in the plasma membrane of the target cell and produce pores with different diameters; these pores alter the permeability of the target plasma membrane to small molecules and even proteins, often leading to cell death.

Innate immune responses combat infectious microbes, but may also drive pathological inflammation. Cell death mechanisms are central for these processes, since they lead to the death of an infected cell, as well as to the release of danger-associated molecular patterns (DAMPs) that drive the inflammatory process in either a beneficial or pathological manner [2]. A cell death mechanism, known as pyroptosis, was shown to be important for controlling several infections by activating the inflammasome complexes and the subsequent secretion of the mature pro-inflammatory cytokines, interleukin 1β (IL-1β) and IL-18 [3,4].

Inflammasomes are multimeric protein complexes that assemble in the cytosol after sensing pathogen-associated molecular patterns (PAMPs) or DAMPs. Their activation is mediated by evolutionarily conserved innate immune pattern recognition receptors (PRRs) [5]. Inflammasomes can be divided into 2 categories: canonical, in which procaspase-1 is converted into a catalytically active enzyme [6], and noncanonical, which are initiated by the activation of caspase-11 [7]. The canonical inflammasome contains a nucleotide-binding oligomerization domain (NOD), leucine-rich repeat (LRR)-containing protein (NLR) family member (e.g., NLRP1, NLRP3, or NLRC4), or the DNA sensor, Absent in Melanoma 2 (AIM2). NLRs and AIM2 contain a pyrin domain (PYD) or a caspase recruitment domain (CARD) [8] that interacts with the apoptosis-associated speck-like protein containing a CARD (ASC) adaptor or with procaspase-1, directly. This interaction leads to dimerization, autocleavage, and activation of caspase-1 [9], which further cleaves the inactive precursors of IL-1βand IL-18 into their active, pro-inflammatory forms, and directs their secretion; it may also lead to pyroptotic cell death [10,11].

We previously discovered a functional and potent pore-forming hemolysin produced by the Gram-negative marine bacterium *Vibrio proteolyticus* (*V. proteolyticus*), termed VPRH [12]. *V. proteolyticus*, which was first isolated from the gut of the wood borer *Limnoria tripunctata* [13], is pathogenic to marine animals; it was isolates as part of a *Vibrio* consortium from yellow band diseased corals [14], and was shown to cause mortality in fish [15] and in the crustacean model organism, *Artemia* [16]. The hemolysin, VPRH, is a 305-residue protein containing a secretion signal peptide followed by a leukocidin domain. Although the VPRH leukocidin domain is homologous to those found in other known PFTs, such as α-hemolysin, HlyA, and LukED, VPRH defines a distinct and understudied class within the leukocidin superfamily [12]. Members of the VPRH leukocidin class are confined to marine bacteria, including emerging pathogens of humans and marine animals. Recently, we showed that when VPRH was introduced to cultures of human epithelial HeLa cells, it caused changes in the actin cytoskeleton, resulting in cell lysis. Similar VPRH-dependent lytic activity was also found when *V. proteolyticus* bacteria were added to murine RAW 264.7 macrophage cell cultures [12].

A common result of PFT insertion into the plasma membrane is a drop in cellular potassium concentration, which leads to activation of signaling cascades such as the inflammasome and mitogen-activated protein kinase pathways [17]. Several pore-forming leukocidins, such as *Staphylococcus aureus* α-hemolysin [18] and Panton-Valentine leukocidin, were found to affect inflammasome activation in mammalian immune cells. Since VPRH was only tested against cells that do not possess a functional inflammasome (HeLa and RAW 264.7), it is not known whether members of the VPRH class of leukocidins affect immune cells similarly.

In this work, we sought to determine whether VPRH affects the inflammasome, and if so, to decipher the underlying mechanism. Importantly, we found that VPRH induced a rapid cell death in bone marrow-derived macrophages (BMDMs), in comparison with the slower cell death induced in HeLa and RAW 264.7 cells that do not contain a functional inflammasome [9]. Using chemical inhibitors, we determined that the cell death in BMDMs comprised two distinct steps: the first, a rapid step, was pyroptosis; while the mechanism underlying the second, a slower step, remains unexplored. Furthermore, we demonstrated that VPRH-induced pyroptosis was dependent on the NLRP3 inflammasome, since NLRP3-deficient BMDMs were protected from the initial, rapid cell death. In agreement with these findings, VPRH led to the specific secretion of the pro-inflammatory cytokine IL-1βin a NLRP3-dependent manner. Therefore, we concluded that VPRH induces cell death in mammalian cells; this cell death is accelerated in primary macrophages by rapid activation of the NLRP3 inflammasome and pyroptosis.

## RESULTS AND DISCUSSION

### Accelerated VPRH-induced cell death in primary macrophages

We previously reported that the pore-forming hemolysin, VPRH, induced actin cytoskeleton rearrangement and lysis in HeLa and RAW 264.7 cells upon infection with *V. proteolyticus* [12]. In this work, we sought to determine the effect of VPRH on primary immune cells. To this end, we repeated our original experiments, but with the addition of BMDMs, a known model for immune response cells, and a first-line defense against foreign invaders [19]. Importantly, BMDMs are also known to contain a full set of cell death mechanisms, including pyroptosis and necroptosis, as opposed to HeLa and RAW 264.7 cells [9,19]. As shown in Figure 1A, infection of HeLa and RAW 264.7 cells, as well as BMDMs resulted in VPRH-mediated induction of cell death, as evident by detection of propidium iodide (PI) uptake by cells (PI enters the cells and binds nucleic acids only upon the loss of membrane integrity). Interestingly, the VPRH-mediated cell death was more rapid in BMDMs, suggesting that they are more sensitive than the previously tested HeLa and RAW 264.7 cells (100% BMDM cell death within 3h, compared to 5h for RAW 264.7 and 8h for HeLa). This result implies that cell death mechanisms that are missing in both the HeLa and RAW 264.7 cell lines may play a role in the rapid VPRH-induced cell death observed upon BMDM infection. Notably, a slower, VPRH-independent cell death was observed only in BMDMs upon infection with the *V. proteolyticus* Δ*vprh* mutant. This could be attributed to the high bacterial load found in the infection media several hours post-infection (Figure 1A-B).

**Figure 1.**
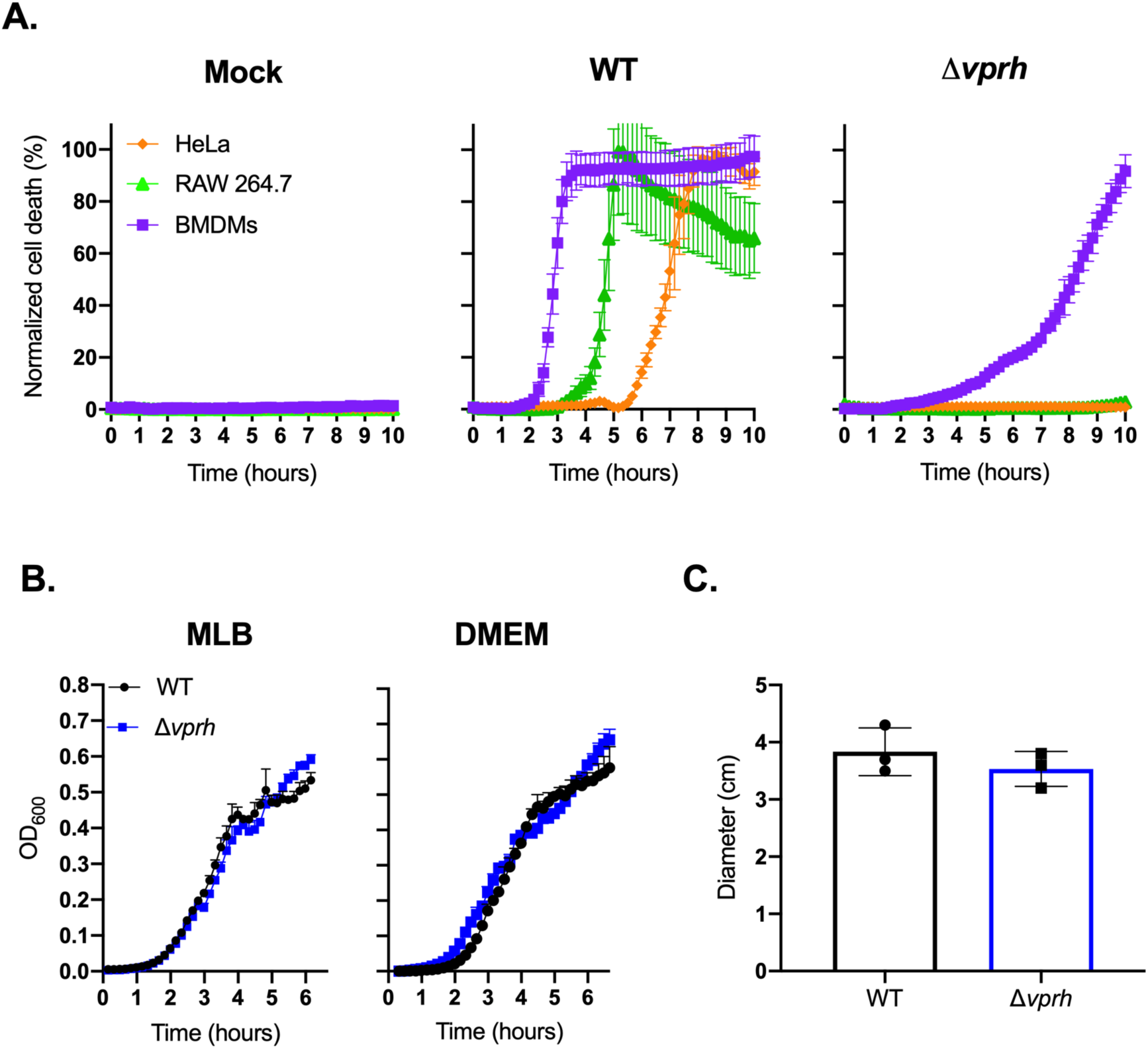
VPRH induces rapid cell death in primary macrophages. **(A)** Assessment of cell death upon infection of mammalian cells with *V. proteolyticus* strains. Approximately 3.5*10^4^ HeLa, RAW 264.7, or BMDM cells were seeded into 96-well plates in triplicate, and were infected with wild-type (WT) *V. proteolyticus* or a mutant in which we deleted *vprh* (Δ*vprh*), at the multiplicity of infection (MOI) 20. Propidium iodide (PI) was added to the medium prior to infection, and its uptake was assessed using real-time microscopy (IncucyteZOOM) and then graphed as the percentage of PI-positive cells normalized to the number of cells in the wells. **(B)** Growth of *V. proteolyticus* strains. Growth of *V. proteolyticus* strains, used in A, MLB, and DMEM media at 30°C, measured as absorbance at 600 nm (OD_600_). **(C)** Motility of *V. proteolyticus* strains. Swimming motility of *V. proteolyticus* strains, used in A, measured as migration on a soft-agar plate after overnight incubation at 30ºC. The data in A, B, and C are presented as the mean ± SD; n=3. Results shown are representative of 3 independent experiments.

The apparent cell death phenotypes in BMDMs were not a result of a fitness difference between the wild-type (WT) *V. proteolyticus* and the Δ*vprh* mutant strain, since we did not detect any difference in growth (either in MLB, the bacterial growth medium, or in DMEM, the medium in which infections were performed) (Figure 1B). Moreover, we did not detect any significant difference in bacterial motility, as determined by the diameter of bacterial migration in a swimming assay (Figure 1C). Taken together, our results indicate that *V. proteolyticus* induces rapid cell death in BMDMs; this cell death is dependent on the pore-forming toxin, VPRH.

### VPRH-induced cell death in BMDMs is contact independent

Bacteria-induced cell death may be either dependent or independent of contact with host cells. To test the hypothesis that VPRH-induced cell death in BMDMs is contact independent, as we have shown previously in RAW 264.7 and HeLa cells, we eliminated any direct contact between the bacteria and BMDMs using two approaches. In the first approach, we infected BMDMs with *V. proteolyticus* strains for 3.5 hours in 6-well culture plates while monitoring PI uptake (to measure cell death). Briefly, supernatants were collected from these wells, filtered (0.22 μm filter) to eliminate live bacteria, and then added to untreated BMDMs seeded in 96-well plates. Unfiltered supernatants from the same wells (containing live bacteria) were used as controls. As shown in Figure 2A, the supernatant of the WT-infected wells was still able to induce BMDM cell death, even after filtration (the black open circles), indicating that death was induced by a secreted protein present in the media.

**Figure 2.**
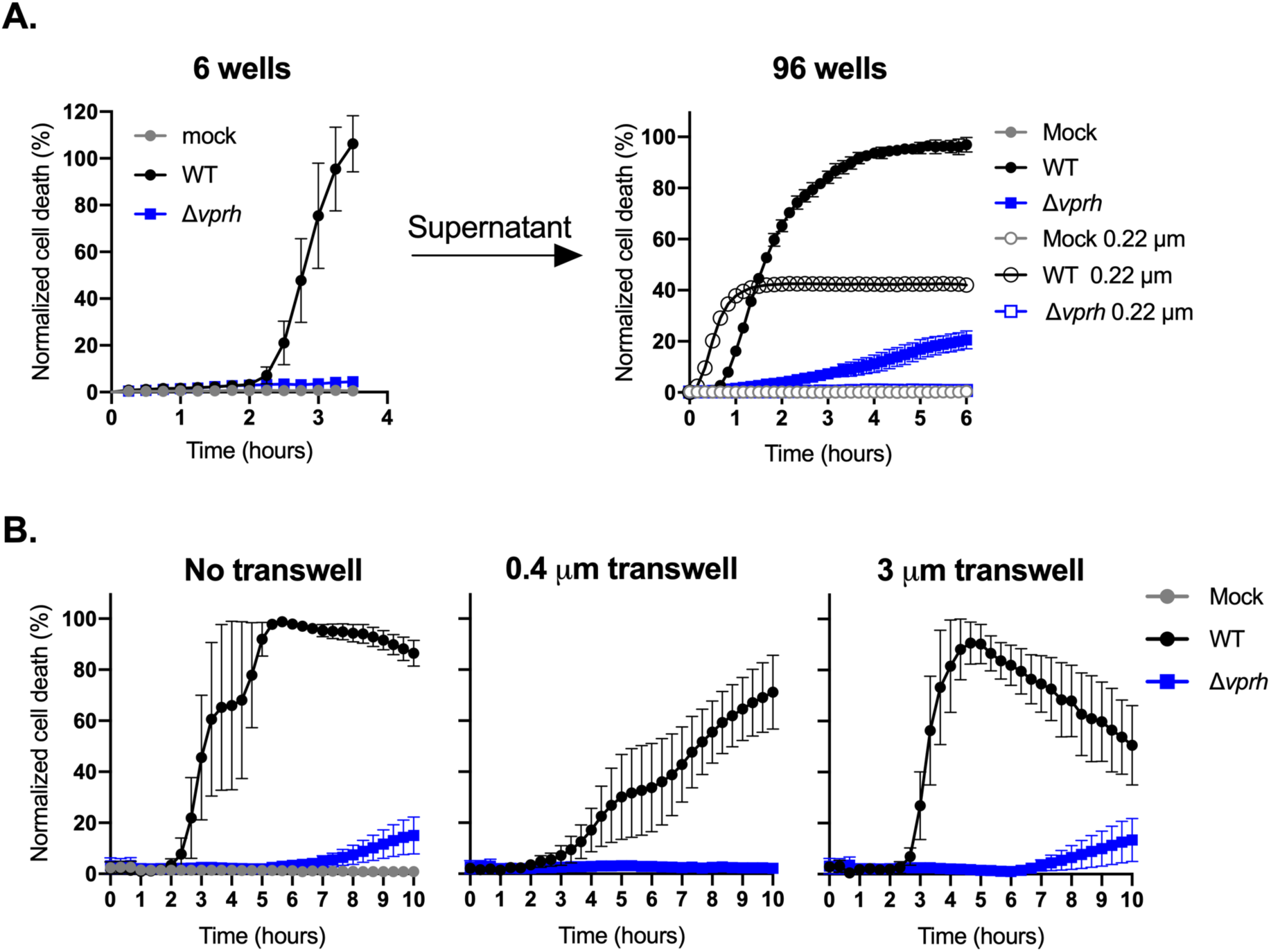
VPRH-mediated cell death is contact independent. **(A)** Approximately 2*10^6^ wild-type BMDMs were seeded into 6-well plates, and were infected with wild-type (WT) *V. proteolyticus* or a Δ*vprh* mutant at MOI 20. Supernatants collected 3.5 hours post-infection were either filtered (0.22 μm filter) or not. Unfiltered or filtered supernatants were added in triplicate to BMDMs in a 96-well plate (3.5*10^4^ BMDMs per well). **(B)** *V. proteolyticus*, as in A, were added to cultures of 10^5^ BMDMs on top of 0.4 μm or 3 μm transwell filters, or without a transwell, in a 24-well plate. In A and B, cell death was determined by monitoring propidium iodide (PI) uptake using real-time microscopy (IncucyteZOOM) and then graphed as the percentage of PI-positive cells normalized to the number of cells in wells. The data are presented as the mean ± SD; n=3. Results shown are representative of 3 independent experiments.

In the second approach, we physically separated the bacteria from the BMDMs using 0.4 μm transwells, and monitored cell death using PI uptake measurements (Figure 2B). Even when physically separated, the bacteria induced VPRH-dependent cell death in BMDMs, as they did when no transwells were used, or when 3 μm transwells (permeable to *V. proteolyticus* bacteria) were used. Taken together, these results confirm that VPRH-mediated cell death is caused by a secreted toxin in a contact-independent manner.

### VPRH induces a rapid, inflammasome-dependent pyroptotic cell death in BMDMs

Since hemolysins were previously shown to induce inflammasome-mediated cell death [20], we hypothesized that the rapid VPRH-mediated cell death in BMDMs was also dependent on inflammasome activation. To test this, we added specific inflammasome inhibitors to BMDM cultures 30 minutes prior to infection with *V. proteolyticus* and monitored their effect on VPRH-induced cell death. The inhibitors used were as follows: i) MCC950, which blocks ASC oligomerization by inhibiting the canonical and non-canonical NLRP3 inflammasome [21]; ii) Vx765, a potent and selective competitive inhibitor of caspase-1 and −4 [22]; and iii) Necrosulfonamide (NSA), a human specific mixed lineage kinase domain-like (MLKL) inhibitor that does not bind to the murine version of MLKL. When applied to murine cells, NSA blocks inflammasome priming, caspase-1 activation, and gasdermin D (GSDMD) pore formation [23]. DMSO was used as the solvent control. As shown in Figure 3A, the addition of NSA or Vx765, both inhibitors of caspase-1 activity, resulted in almost a two-hour delay of the VPRH-mediated cell death; the addition of MCC950, which specifically inhibits the NLRP3 inflammasome, resulted in a shorter, one-hour delay of cell death. Taking the effects of these three independent, inflammasome pathway inhibitors together, we hypothesized that VPRH induces a rapid cell death in BMDMs via the inflammasome-dependent pyroptotic pathway. Since a delayed cell death was still evident, even in the presence of the inflammasome inhibitors, we also hypothesized that a second step, which is inflammasome independent, plays a role in the VPRH-induced cell death in BMDMs.

**Figure 3.**
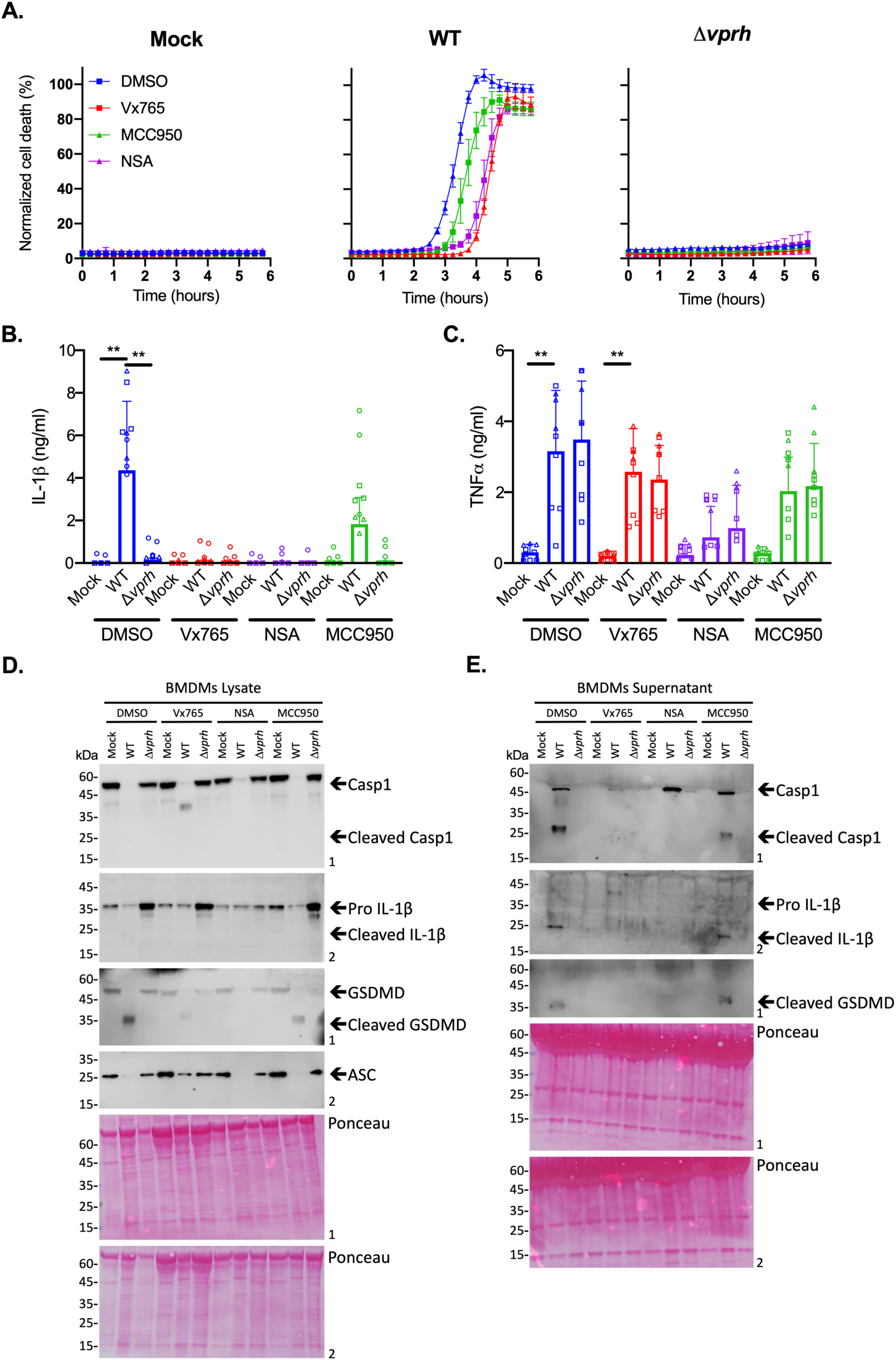
Inflammasome inhibitors delay VPRH-mediated cell death. Approximately 3.5*10^4^ wild-type BMDMs were seeded into 96-well plates in triplicate, and were primed using LPS (100 ng/ml) for 3 hours prior to infection with *V. proteolyticus* mutants at MOI 50. Where indicated, inflammasome inhibitors, Vx765 (25 μM), MCC950 (20 μM), and NSA (20 μM), with the addition of propidium iodide (PI) (1 μg/ml), were added to the cells 30 minutes prior to bacterial infection. DMSO was added as the solvent control. **(A)** PI uptake was assessed using real-time microscopy (IncucyteZOOM) and then graphed as the percentage of PI-positive cells normalized to the number of cells in wells. Data are presented as the mean ± SD; n=3. **(B-E)** Cell lysates and supernatants from experiments described in A were collected 6 hours post-infection. **(B, C)** IL-1β (B) and TNFα (C) secretion were measured using commercial ELISA kits. Statistical comparisons between the different bacterial mutants were carried out for each inhibitor treatment using RM one-way ANOVA, followed by Sidak’s multiple comparison test; the symbols represent 3 independent biological experiments performed in triplicate; the results are presented as the mean (bars) ± SD of 3 independent experiments); ** *P<0.01*. **(D, E)** Inflammasome components, caspase-1 (Casp1), GSDMD, and IL-1β cleavage were detected in BMDM lysates (D) and supernatants (E) using immunoblots (the numbers on the right of each blot indicate the blot number). The data in A, D, and E are representative of 3 independent experiments.

To further support our hypothesis that VPRH induces a rapid, inflammasome-mediated cell death in BMDMs, we sought to determine whether the inflammasome-dependent cytokine, IL-1β, was secreted upon infection with *V. proteolyticus*. To this end, we determined the amount of the cytokines IL-1β and TNFα (an NF-κB-dependent, inflammasome-independent cytokine that was used as a control) in the supernatants of the infected cultures described above (Figure 3A). In agreement with the above results, the addition of inflammasome inhibitors eliminated IL-1β secretion (Figure 3B), whereas they had no effect on TNFα secretion (Figure 3C). These results confirm that VPRH induces an inflammasome-dependent cell death in BMDMs.

We also determined the effect that inflammasome inhibitors had on VPRH-mediated, inflammasome-dependent cell death by monitoring the cleavage and release of caspase-1, IL-1β, and GSDMD in BMDMs infected with WT *V. proteolyticus*. As shown in Figure 3D-E, Vx765 and NSA, which resulted in a two-hour delay in cell death, also inhibited the various inflammasome-dependent phenotypes that were tested: i) caspase-1 processing and release to the supernatant, ii) IL-1β maturation and secretion, and iii) GSDMD processing. Notably, MCC950, which had a milder effect on cell death, was also less effective in inhibiting these inflammasome-dependent phenotypes. Collectively, these findings suggest that the rapid cell death apparent in BMDMs is mediated by VPRH-induced activation of pyroptosis, possibly via the NLRP3 inflammasome.

### VPRH activates the NLRP3 inflammasome in BMDMs

Inflammasome activation may be induced by several NLR family members, including NLRP1 and NLRP3 [24]. The most studied inflammasome is NLRP3, which was shown to be activated by many DAMPs and PAMPs. Furthermore, NLRP3 was also shown to be activated by the necroptotic cell death pathway, which is induced by pseudo-kinase MLKL [25,26]. Thus, to identify the specific inflammasome pathway that was activated by VPRH in BMDMs, we cultured BMDMs from knockout (KO) mice in which different inflammasome activation pathways were severed (i.e., *Mlkl*^−/−^, *Nlrp1*^−/−^, and *Nlrp3*^−/−^). We used real-time microscopy to compare their cell death kinetics to those of BMDMs from WT mice after *V. proteolyticus* infection. We also monitored IL-1βsecretion using ELISA, as well as by processing various inflammasome components to determine inflammasome activation in BMDMs from the KO mice.

As shown in Figure 4A, infection of BMDMs from *Nlrp1*^−/−^ and *Mlkl*^−/−^ mice resulted in rapid cell death, with kinetics comparable to those observed in BMDMs from WT (B6J) mice. Remarkably, infection of *Nlrp3*^−/−^ BMDMs resulted in a delayed cell death phenotype (Figure 4A), similar to the phenotype observed when inflammasome inhibitors were added to WT BMDMs (Figure 3A). These results indicate that NLRP3, but not NLRP1 or MLKL, is required for the rapid cell death induced by VPRH in BMDMs. In agreement with these results, IL-1β secretion was abrogated specifically in *Nlrp3*^−/−^ BMDMs (Figure 4B). In contrast, TNFα secretion was not affected in either of the KO mouse BMDMs (Figure 4C), thus confirming that priming (i.e., the NF-κB-dependent pathway) remained unaffected in the *Nlrp3*^−/−^ BMDMs. In further support of the above results and conclusions, cleaved caspase-1, mature IL-1β, and cleaved GSDMD were absent in *Nlrp3*^−/−^ BMDM culture lysates and supernatants (Figure 4D-E).

**Figure 4.**
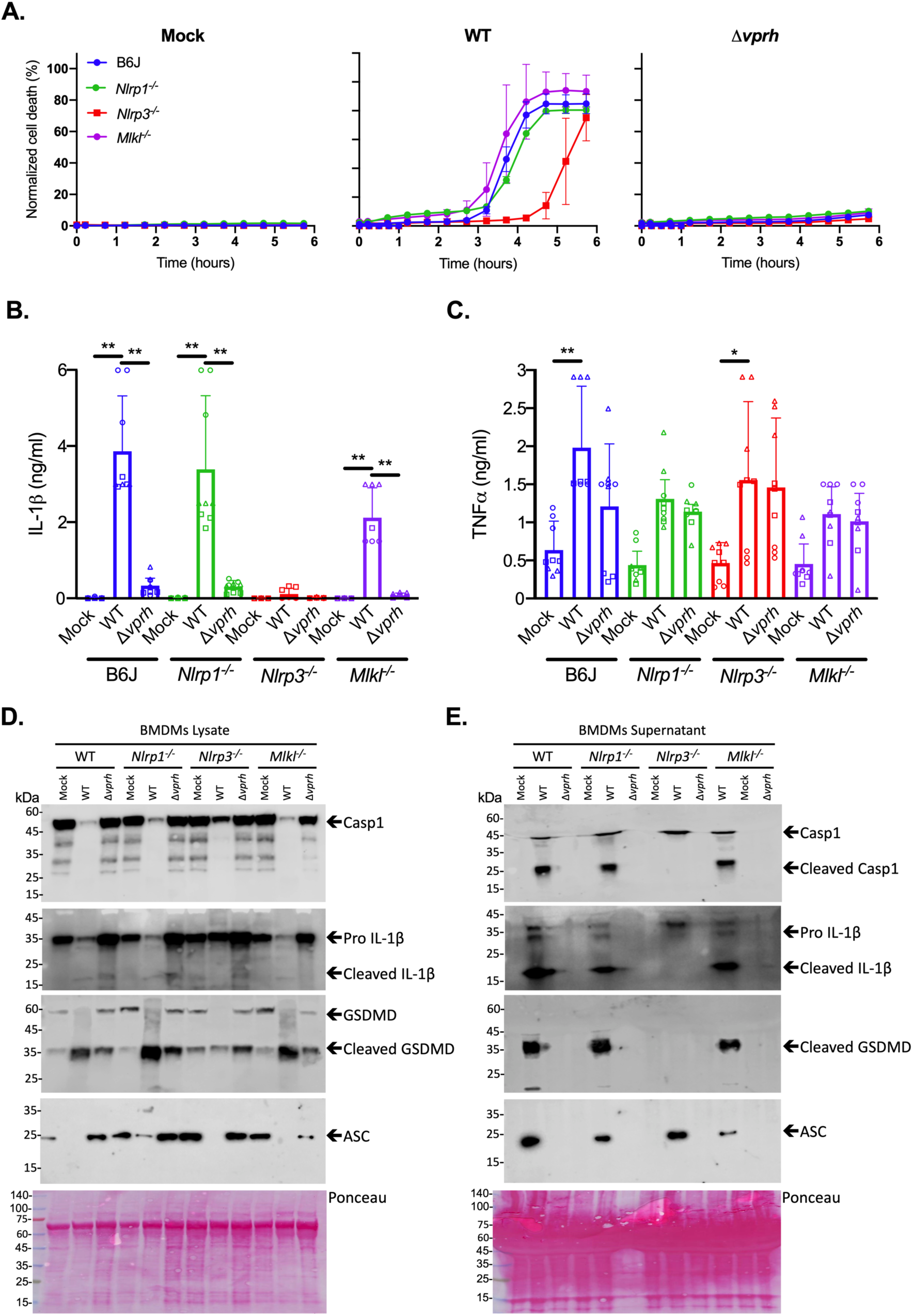
VPRH induces cell death via the NLRP3 inflammasome. Approximately 3.5*10^4^ wild-type (B6J), *Nlrp1*^−/−^, *Nlrp3*^−/−^, and *Mlkl*^−/−^ BMDMs were seeded into 96-well plates in triplicate, and were primed using LPS (100 ng/ml) for 3 hours prior to infection with *V. proteolyticus* mutants at MOI 20. **(A)** PI uptake was assessed using real-time microscopy (IncucyteZOOM) and then graphed as the percentage of PI-positive cells normalized to the number of cells in wells. Data are presented as the mean ± SD; n=3. **(B-E)** Cell lysates and supernatants from experiments described in A were collected 6 hours post-infection. **(B, C)** IL-1β (B) and TNFα (C) secretion were measured using commercial ELISA kits. Statistical comparisons between the different bacterial mutants were carried out for each BMDM genotype using RM one-way ANOVA, followed by Sidak’s multiple comparison test; the symbols represent 3 independent biological experiments performed in triplicate; the results are presented as the mean (bars) ± SD of 3 independent experiments); * *P<0.05*, ** *P<0.01*. **(D, E)** Inflammasome components and caspase-1 (Casp1), GSDMD, and IL-1β cleavage were detected in BMDM lysates (D) and supernatants (E) using immunoblots (the numbers on the right of each blot indicate the blot number). The data in A, D, and E are representative of 3 independent experiments).

### Exogenous complementation of VPRH restores NLRP3-dependent cell death in BMDMs

To further confirm our hypothesis that VPRH induces NLRP3 inflammasome-dependent cell death in BMDMs, we introduced *vprh* on an arabinose-inducible expression plasmid (pVPRH) back into the Δ*vprh* strain. As shown in Figure 5, complementation of VPRH restored all of the inflammasome-mediated phenotypes upon BMDM infection, including rapid cell death (Figure 5A), IL-1β secretion (Figure 5B), as well as cleavage of caspase-1, maturation of IL-1β, and cleavage of GSDMD (Figure 5D-E). Notably, exogenous expression of VPRH had no effect on TNFα secretion (Figure 5C). Importantly, the phenotypes that were observed with the VPRH-complemented strain were NLRP3-dependent, as evident by the fact that none of them were restored when *Nlrp3*^−/−^ BMDMs were infected (Figure 5).

**Figure 5.**
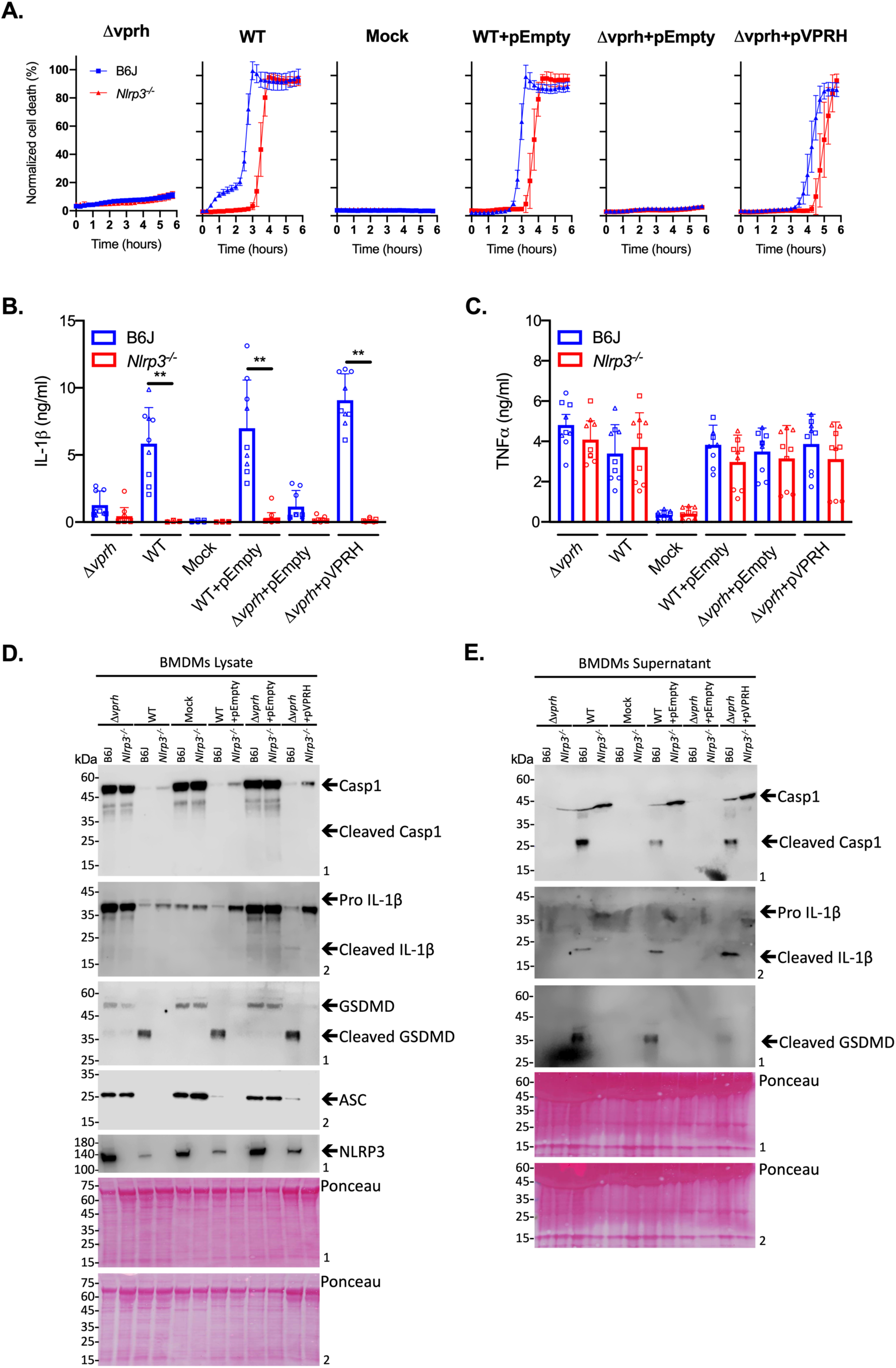
Exogenous complementation of VPRH restores the NLRP3 inflammasome activation in BMDMs. Approximately 3.5*10^4^ wild-type (B6J) and *Nlrp3^−/^*^−^ BMDMs were seeded into 96-well plates and then primed using LPS (100 ng/ml) for 3 hours prior to infection with *V. proteolyticus* mutants at MOI 50. **(A)** PI uptake was assessed using real-time microscopy (IncucyteZOOM) and graphed as the percentage of PI-positive cells normalized to the number of cells in the wells. Data are presented as the mean ± SD; n=3. **(B-E)** Cell lysates and supernatants from experiments described in A were collected 6 hours post-infection. **(B, C)** IL-1β (B) and TNFα (C) secretion were measured using commercial ELISA kits. Statistical comparisons between the different BMDMs were carried out for each bacterial mutant using RM one-way ANOVA, followed by Sidak’s multiple comparison test; the symbols represent 3 independent biological experiments performed in triplicate; the results are presented as the mean (bars) ± SD of 3 independent experiments); ** *P<0.01*. **(D, E)** Inflammasome components, caspase-1 (Casp1), GSDMD, and IL-1β cleavage were detected in BMDM lysates (D) and supernatants (E) using immunoblots (the numbers on the right of each blot indicate the blot number). The data in A, D, and E are representative of 3 independent experiments).

## CONCLUDING REMARKS

We previously showed that the *V. proteolyticus* VPRH, a leukocidin domain-containing hemolysin, induces cell death upon infection of HeLa and RAW 264.7 cells [12]. Here, we found that this cell death phenotype is accelerated when BMDMs are infected. Since BMDMs, unlike HeLa and RAW 264.7 cells, possess functional inflammasomes whose activation can lead to rapid cell death, we hypothesized that VPRH activates the inflammasome in BMDMs. Our results clearly indicate that in BMDMs, VPRH induces a two-step cell death. Although the mechanism leading to the second, slower step remains to be elucidated, we showed that the first, rapid step is mediated by VPRH-dependent activation of the inflammasome. It is plausible that the second, slower step of cell death observed in BMDMs is mediated by the same mechanism that induced VPRH-dependent cell death in HeLa and RAW 264.7 cells that lack a functional inflammasome. Using a combination of genetic and chemical approaches, we identified the NLRP3 inflammasome as the specific pathway responsible for the VPRH-induced rapid cell death in BMDMs. Taken together, our results shed new light on the virulence potential of VPRH, and possibly that of other members of this leukocidin class of pore-forming toxins. Although VPRH induces cell death in non-immune cells, such as HeLa cells, it can also specifically activate innate immune response mechanisms in primary macrophages by activating the NLRP3-inflammasome pathway. It remains to be investigated whether activation of NLRP3 inflammasome-mediated cell death benefits the pathogen or the host during infection.

## METHODS

### Reagents

Unless stated otherwise, all cell culture reagents were purchased from Biological Industries, Beit-Haemek, Israel. Lipopolysaccharides (LPS) and propidium Iodide (PI) were purchased from Sigma-Aldrich. Necrosulfinamide (NSA) was purchased from Tocris Bioscience; Vx765 and MCC950 were purchased from Invitrogen. HRP-conjugated secondary antibodies were purchased from Jackson ImmunoResearch Labs (West Grove, PA, USA). ELISA kits were purchased from eBioscience or R&D.

### Mice

C57/BL6/J (wild-type [B6J]), Nlrp3A350VneoR/+, which are NLRP3 KO, NLRP1 KO, and MLKL KO mice, were bred under specific pathogen-free conditions in the animal facility at Tel Aviv University. Experiments were performed according to the guidelines of the Institutes’ Animal Ethics Committees.

### Cell culture

HeLa, RAW 264.7, and BMDM cells were grown in DMEM culture medium containing 10% FBS, 1% penicillin-streptomycin, and 1% HEPES, at 37°C, in a 5% CO2 incubator.

### Bone marrow-derived macrophages

Bone marrow (BM) cells from mice were isolated by flushing femurs and tibias with 5 ml PBS, supplemented with 2% heat-inactivated fetal bovine serum (FBS) Gibco (Thermo Fisher Scientific, Waltham, MA, USA). The BM cells were centrifuged once and then re-suspended in tris-ammonium chloride at 37°C for 30 seconds to lyse red blood cells. The cells were centrifuged again and then strained through a 70 μm filter before being re-suspended in DMEM supplemented with 10% FBS. BMDMs were obtained by 7 days differentiation with L-con media as previously described [27].

### Bacterial strains and media

*Vibrio proteolyticus* strain ATCC 15338 (also termed NBRC 13287) and its derivatives were routinely grown in Marine Lysogeny Broth (MLB; Lysogeny broth supplemented with NaCl to a final concentration of 3% w/v) at 30°C. To induce the expression of genes from a plasmid, 0.1% (w/v) L-arabinose was included in the media. When necessary to maintain plasmids, the media were supplemented with 250 µg/ml kanamycin. Construction of the *vprh* (locus tag VPR01S_RS09275; encoding WP_021705060.1) deletion strain (Δ*vprh*) and of the VPRH arabinose-inducible expression plasmid (pVPRH) were reported previously [12].

### Bacterial growth assay

Overnight-grown cultures of *V. proteolyticus* were normalized to an OD_600_=0.01 in MLB media and transferred to 96-well plates (200 µl per well). For each experiment, n=3. Cultures were grown at 30°C in a BioTek EPOCH2 microplate reader with continuous shaking at 205 cpm. OD_600_ readings were acquired every 10 minutes. Experiments were performed at least three times with similar results.

### Bacterial swimming assay

Swimming media plates were prepared with Lysogeny broth containing 20 g/l NaCl and 3 g/l Agar. *V. proteolyticus* strains that were grown overnight on a MLB plate were picked and then stabbed into the swimming plates using a toothpick (n=3). Plates were incubated at 30°C for 8-16 hours. Swimming was assessed by measuring the diameter of the spreading bacteria. Experiments were performed at least three times with similar results.

### Infection of cell cultures

Unless otherwise stated, HeLa, RAW 264.7, and BMDM cells were washed 3 times using PBS and seeded in a final concentration of 1.75*10^5^/ml in triplicate in 1% FBS and penicillin-streptomycin-free DMEM media. Unless otherwise stated, cells were infected with *V. proteolyticus* mutants at MOI 20 or 50, as indicated. Where indicated, BMDMs were pre-incubated with LPS (100 ng/ml, 3 hours). When used, inflammasome inhibitors Vx765 (25 μM), MCC950 (20 μM), and NSA (20 μM) were added 30 minutes prior to infection. More specifically, overnight cultures of *V. proteolyticus* mutants were diluted and prepared in DMEM without antibiotics. Bacteria were added to the wells at the final MOI 20, and plates were centrifuged for 5 minutes at 400Xg. Plates were then inserted into IncucyteZOOM for incubation and for monitoring cell death.

### Live cell imaging

Plates with infected cells were placed in IncucyteZOOM (Essen BioScience) and images were recorded every 10-30 minutes. Data were analyzed using IncucyteZoom2016B analysis software and exported to GraphPad Prism software. Normalization was then performed according to the maximal PI-positive object count to calculate the percentage of dead cells.

### Immunoblot analyses of proteins

Cells were collected and centrifuged for five minutes at 400xg (4°C) in order to separate them from the supernatant. Next, the cells were lysed either by using RIPA buffer in the presence of protease inhibitors at 4°C for 15 minutes, or directly by applying buffer from denaturing western blot samples to cells. Lysed cells were loaded onto any kD gradient Criterion TGX-Free precast gels (Bio-Rad). Proteins were transferred onto a nitrocellulose membrane (Bio-Rad), and Ponceau S staining was performed routinely to evaluate the loading accuracy. Membranes were blocked with 5% (w/v) skim milk in TBS for 1-2 hours, and then probed overnight with primary antibodies (all diluted 1:1000, unless noted otherwise): mouse-NLRP3 (AdipoGen; cryo-2), mouse-ASC (AdipoGen; AL177), pro and mature mouse-IL-1β(R&D Systems; AF-401-NA), pro and cleaved mouse caspase-1 (Santa Cruz; sc-514) (Adipogen; AG-20B-0042-C100), and pro and cleaved mouse-GSDMD (Abcam; ab209845). Relevant horseradish peroxidase-conjugated secondary antibodies were applied for at least 1 hour. Membranes were washed four times in TBS containing 0.1% (v/v) Tween 20 (TBST) between antibody incubations. Antibodies were diluted in TBST containing 5% skim milk. Immunoblots were developed using an ECL kit (Bio-Rad) in an ODYSSEY Fc (Li-COR) equipped with Image Lab software. All images were cropped for presentation; their full-size images will be presented upon request.

### Statistics

Unless otherwise specified, data are presented as the mean ± standard deviation (SD). Comparisons were performed using RM one-way ANOVA, followed by Sidak’s multiple comparison test. For each test, *P* values <0.05 were considered statistically significant.

## ACKNOWLEDGMENTS

This work was performed in partial fulfillment of the requirements for a Master’s degree for (N.B.) and a Ph.D. degree for (H.C.), the Sackler Faculty of Medicine, Tel Aviv University, Israel. We would also like to thank Chaya Mushka Fridman, Rotem Ben-Yaakov, and Yasmin Dar for their technical assistance in maintaining and growing the *V. proteolyticus* strains, and Ziv Erlich in generating the BMDMs. The research of M.G. was supported by the Israel Science Foundation (ISF) (grant #1416/15 and 818/18), the Margot Stoltz Foundation through the Faculty of Medicine grants of Tel-Aviv University, and individual research grants from the Varda and Boaz Dotan Research Center. The research of D.S. was supported by the ISF (grant #920/17).

## AUTHOR CONTRIBUTIONS

Conceptualization, D.S. and M.G.; Methodology, N.B., H.C., L.E-B., D.S., and M.G.; Investigation, N.B., H.C., L.E-B., D.S., and M.G.; Writing – Original Draft, D.S. and M.G.; Writing – Review & Editing, N.B., H.C., L.E-B., D.S., and M.G.; Funding Acquisition, D.S. and M.G.; Validation, L.E-B., H.C.; Supervision, D.S. and M.G.

## COMPETING INTERESTS

The authors declare no competing interests.

